# Mitonuclear interactions affect locomotor activity and sleep in *Drosophila melanogaster*

**DOI:** 10.1101/2021.10.30.464953

**Authors:** Lucy Anderson, M. Florencia Camus, Katy M. Monteith, Tiina S. Salminen, Pedro F. Vale

## Abstract

Mitochondria are organelles that produce cellular energy in the form of ATP through oxidative phosphorylation, and this primary function is conserved between many taxa. Locomotion is a trait that is highly reliant on metabolic function and expected to be greatly affected by disruptions to mitochondrial performance. To this end, we aimed to examine how activity and sleep vary between *Drosophila melanogaster* strains with different geographic origins, how these patterns are affected by mitochondrial DNA (mtDNA) variation, and how breaking up co-evolved mito-nuclear gene combinations affect the studied activity traits. The results demonstrate that *Drosophila* strains from different locations differ in sleep and activity, and the extent of variation differs between sexes, females in general being more active. By comparing activity and sleep of mtDNA variants introgressed onto a common nuclear background in cytoplasmic hybrid (cybrid) strains, we establish that mtDNA variation affects both traits, sex specifically. Furthermore, by using previously published mtDNA copy number data, we detected a positive correlation between mtDNA copy number and the activity levels of the cybrid flies. Altogether, our study shows that both mtDNA variation and mitonuclear interactions affect activity and sleep patterns, highlighting the important role that both genomes play on life-history trait evolution.

## Introduction

Mitochondria are key organelles in a range of critical metabolic processes and are the primary energy producers for the eukaryotic cell. In addition to this primary role, they are also involved in a range of other vital processes that control key aspects of cellular growth and regulation such as signalling (1), cellular differentiation (2), cell death (3) and immunity (4). The mitochondrial machinery responsible for ATP production via oxidative phosphorylation (OXPHOS) is jointly encoded by the mitochondrial (mtDNA) and nuclear genomes. All the thirteen protein coding genes of the mtDNA genome are involved with OXPHOS, as majority of the OXPHOS subunits as well as over 1200 other genes required for mitochondrial functions are encoded by nuclear DNA (5-8). In addition to vast differences in genome size, there are large differences in genome copies, with hundreds of mtDNA copies inhabiting each diploid cell (9). Consequently, precise and synchronised coordination between the two genomes is required for proper assembly and function of the components of the electron transport chain and mitochondrial functions. Disruptions to this system – by mutations in the mitochondrial or nuclear counterparts – can have consequences to a wide-range of life-history phenotypes (10), and in severe cases leads to mitochondrial disease (11-15).

A consequence of this tight intergenomic partnership is that any trait with heavy metabolic underpinnings is reliant on the compatibility between mitochondrial and nuclear genomes. Two such traits which are vital for everyday function are locomotor activity and sleep. Sleep is integral to regular brain function, influencing processes such as learning and memory (16), and plays a role in cellular processes such as metabolic recovery and oxidative stress (17). Continued sleep deprivation results in fatality for both invertebrate and vertebrate species (18-20). In humans, sleep deprivation is associated with an increased risk of metabolic and cognitive disorders (21). Locomotion is also heavily reliant on metabolic function, with studies finding a strong positive correlation between activity and resting metabolic rate (22). Moreover, deviations of optimal organismal activity as well as exercise intolerance have been linked to several metabolic disorders originating from mitochondrial dysfunction (23). For instance, *Drosophila* models of neuromuscular degeneration have shown a progressive decrease in fly activity with time (24). Additionally, nutritional metabolic interventions in the form of dietary restriction have shown to change activity patterns (25).

Given their tight link with metabolism, mutations affecting mitochondrial function are predicted to affect both sleep and locomotor activity. A previous review of primary mitochondrial diseases described sleep disorders in humans are associated with mutations in mitochondrial DNA (26). However, there are currently few clear causal links between specific mitochondrial mutations and disrupted activity and sleep patterns (26, 27). The fruit fly, *Drosophila melanogaster*, offers a powerful system to address the link between mitochondrial variation and activity and sleep disruption. *Drosophila* is an established genetic model system, including the study of mito-nuclear effects on various phenotypic traits (28-31). As mtDNA is maternally inherited, introgression enables the generation of flies with specific combinations of nDNA and mtDNA. Generation of cytoplasmic hybrid (cybrid) lines, in which flies contain a mtDNA variant on a controlled nuclear background, allows the effects of mtDNA mutations disentangled from nuclear genome variation (32). A large body of work using *Drosophila* cybrid lines has established that several measures of life-history phenotypes are modulated by changes of the mitochondrial genome, including aging (33-35), fitness (31, 36, 37), and metabolic rate (38).

*Drosophila* is also an established model for the study of sleep and circadian rhythms and display a state of quiescence which shares critical features of mammalian sleep (39). These similarities include an elevated arousal threshold (39), altered brain electrical activity (40) and a decrease in amount of sleep as flies age (41). Furthermore, gene expression associated with ‘waking’ in fruit flies has been shown to correlate with ‘waking’ genes in mammals (39). A relevant example is the mtDNA-encoded Cytochrome oxidase C, subunit I, which has been demonstrated to have elevated expression during the initial hours following sleep in both *Drosophila* and rats (39). This homology between *Drosophila* and mammalian sleep, combined with the knowledge that many of the genetic and molecular regulators of sleep are conserved between flies and humans (42), has prompted extensive use of the fruit fly as a genetically tractable model organism in the study of sleep.

To specifically address the role of variation in the mitochondrial DNA on activity and sleep, we examined the sleep-wake cycles and activity profiles of a worldwide collection of eight *D. melanogaster* lines, in addition to a set of derived cybrid lines which contained each of the eight mtDNA variants introgressed onto a single common nuclear background. This experimental setup allowed us to investigate the baseline activity and sleep profiles of each line and to assess the contribution of mtDNA to these phenotypes. Further, because the mitochondrial genome of each line has been previously characterized (37) and presents unique mtDNA variation at the haplotype level as well as common variants at the haplogroup level, we were able to associate this variation on sleep and activity patterns, sex specifically.

## Materials and methods

### Fly strains, backcrossing and rearing conditions

We sourced eight wild-type *D. melanogaster* strains, with distinct geographic origins (Table 1), originally obtained from the Drosophila Stock Center (Bloomington, IN). Cybrid lines were created earlier by backcrossing females from each strain (carrying unique mtDNA variant) to males from the nuclear-donor strain of the Oregon RT strain (Oregon R strain maintained long-term in Tampere, Finland; ORT) for at least 12 generations (37). This resulted in a total of 16 strains; 8 of which were the Bloomington derived strains representing coevolved mito-nuclear combinations and 8 were nORT mtDNAx cybrids. All lines were cultured on standard Lewis medium (43), supplemented with yeast, under 12: 12 light: dark cycles at 25° C and 60% humidity. Flies were propagated by placing 2–4-day old females and males on food vials for 3-4 days, with adults being discarded and egg clutched kept. This rearing regime maintained egg densities low enough to prevent larval overcrowding.

**Table 1.**
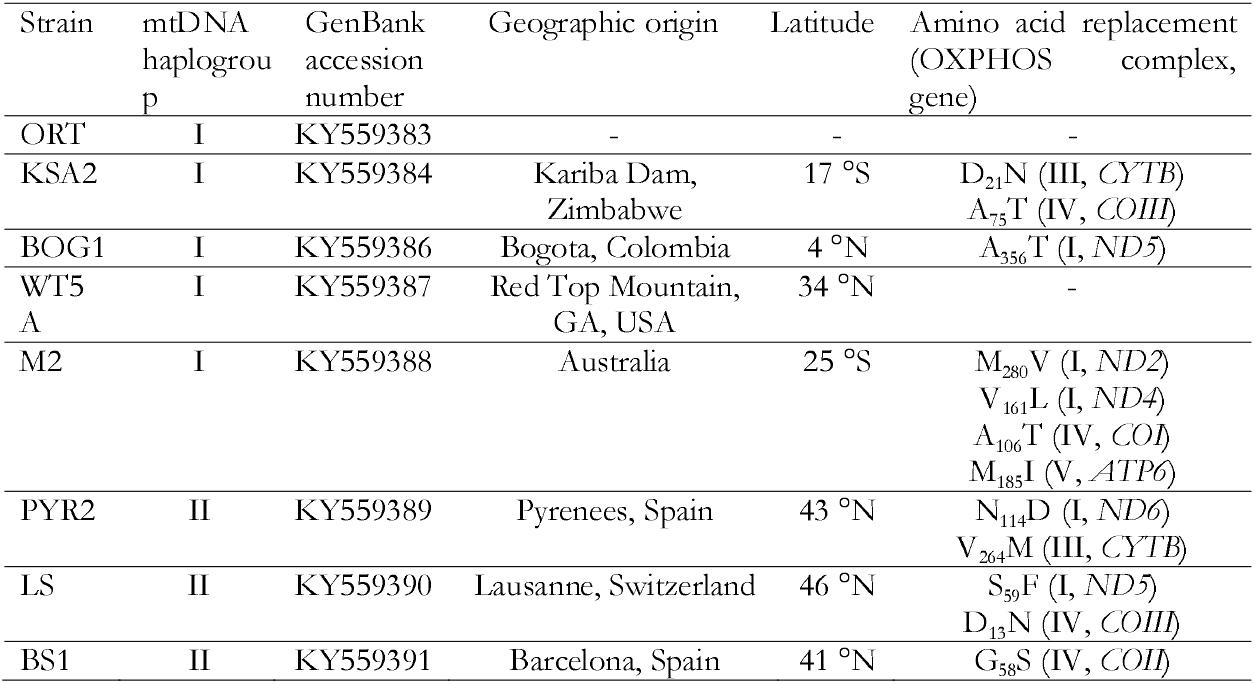
Drosophila melanogaster strains used in the study. Strains were obtained from Bloomington stock centre and originally collected from different continents. GenBank accession numbers refer to the mtDNA coding region sequences. Haplotype-specific amino acid replacements have been added here based on Salminen et al. 2017.

### Experimental design

Tubes for the Drosophila Activity Monitor were prepared prior to the experiments. A solution of 16% sucrose: 2% agar was prepared in distilled water and autoclaved for sterilisation and stored at room temperature before re-melting for use. Each clean class tube (Trikinetics, Waltham, MA) contained approximately 1cm sucrose-agar medium and a rubber cap in one end of the tube and after the anesthetised fly had been inserted into the tube it was closed with a rubber cap containing a small hole for ventilation (44).

Flies from each *coevolved* and *cybrid* strain were collected upon eclosion and kept in food vials for 2 days to make sure flies entering the experiment were mated. Flies were then anesthetised with CO_2_, sorted by sex and transferred into an activity tube using a fine paintbrush. Fifteen individuals per sex and strain were placed in the tubes and randomly positioned in the activity monitors. Each monitor had a null or blank control in which a recorded position contained an empty tube or no tube, respectively. Monitoring of fly activity and sleep lasted for three adjacent days. The assay was split across five experimental blocks, to allow the inclusion of a subset of replicates for each line in all blocks. Each tube was bisected by an infrared beam and locomotor activity movement was recorded whenever a fly broke the beam. The number of activity counts (beam breaks) per minute was stored. In order to quantify sleep from the activity data generated by the DAM, sleep was defined as a 5 minute time-bin with no registered activity (45). To determine whether individuals/flies are more active simply because they sleep less, we calculated the proportion of time each replicate spends sleeping, as well as the mean activity count whilst awake. The experiments were run under 12: 12 light: dark cycles at 25° C at constant temperature and humidity.

### Statistical analysis

Total activity count, mean awake activity and proportion of time sleeping were calculated for each individual fly (see also (46)). Any fly that died during the experiment was excluded from analysis. We analyzed the co-evolved and cybrid lines separately, using otherwise identical statistical models. Total activity count and mean awake activity were analysed using linear mixed effects models (LME). Models fitted ‘line’, ‘sex’ and their interaction as categorical fixed effects. Variation in proportion of time asleep was analysed using generalized linear effects models (GLME) assuming binomial distributed error, which also fitted ‘line’, ‘sex’ and their interaction as categorical fixed effects. All models included the random effect of ‘replicate’ nested within ‘block’ to account for variation between individuals among different blocks. We also investigated if breaking up co-evolved mito-nuclear gene complexes affected each of these behavioral outputs. This analysis therefore included the co-evolved and cybrid fly lines to analyse the effect of mtDNA variant on its original (coevolved) or new (cybrid) nuclear background. Similar model structure was used as described above for each response variable, where each model fitted ‘type’ (coevolved or cybrid), ‘sex’ and their interactions as categorical fixed effects, and ‘line’ as a random effect nested within type.

mtDNA copy number data of three-day old females and males, measured from the whole adult extracts by Salminen et al. (2017) was used to test the correlation between copy number and activity and sleep patterns. mtDNA copy number was originally measured with qPCR by measuring the ratio between nuclear and mitochondrial target genes (37). Nonparametric Spearman’s rank correlation was calculated based on the average mtDNA copy number values and the averages of total activity levels, activity when awake and the proportion flies spent sleeping. Spearman’s rank correlation between mtDNA copy number and the studied activity traits were measured in each sex for either the co-evolved or the cybrid lines. R version 1.1.4 (47) was used for analysis and plots, using packages *ggplot2* (48), *dplyr* (49), lme4 (50), car (51) and plotrix (52). All datasets and full R code for all analyses can be found at https://doi.org/10.5281/zenodo.5573904.

## Results

### Co-evolved fly lines show sex-specific natural variation in sleep and activity patterns

We first evaluated the sleep and activity patterns of genetically and geographically diverse fly lines carrying coevolved mito-nuclear combinations (Table 1). Activity profiles showed that all lines were crepuscular, exhibiting a peak of activity at the onset of the dark period (Fig S1). In the majority of strains, females were significantly more active than males, although the extent of this difference was influenced by genotype and in lines BS1 and WT5A, males were more active than females (Table 2 ‘Line x Sex’ effect; Figure 1A). Overall, males spent a higher proportion of time asleep compared to females, and the extent of this difference varied between genetic backgrounds (Table 2 ‘Line x Sex’ effect; Figure 1B-C). Notably, while males slept for a greater proportion of the day, they were significantly more active while awake compared to females, and again the extent of this variation differed between lines (Table 2 ‘Line x Sex’ effect; Figures 1B-C). This suggests that although females exhibit higher total activity, this is due to females spending more time awake (sleeping less) rather than having higher levels of activity when awake.

**Table 2.** Summaries of test statistics for fixed effects in linear models. See methods for model details.

|  | *Total Activity Count* |  | *Proportion asleep* |  | *Mean awake activity* |  |
| --- | --- | --- | --- | --- | --- | --- |
| | $\chi^2$ | P | $\chi^2$ | P | $\chi^2$ | P |
| ***Original Lines*** |  |  |  |  |  |  |
| *Line* | 320.74 | < 0.001 | 12808.10 | < 0.001 | 10.56 | 0.159 |
| *Sex* | 3.79 | 0.052 | 1063.30 | < 0.001 | 9.40 | 0.002 |
| *Line* × *Sex* | 59.90 | < 0.001 | 1943.80 | < 0.001 | 16.48 | 0.021 |
| ***Cybrid lines*** |  |  |  |  |  |  |
| *Line* | 40.80 | < 0.001 | 2563.42 | < 0.001 | 47.32 | < 0.001 |
| *Sex* | 134.56 | < 0.001 | 5683.25 | < 0.001 | 45.47 | < 0.001 |
| *Line* × *Sex* | 21.95 | 0.003 | 904.27 | < 0.001 | 20.11 | 0.005 |
| ***Coevolved vs. Novel*** |  |  |  |  |  |  |
| *Type* | 18.62 | < 0.001 | 10.42 | 0.0012 | 8.22 | < 0.001 |
| *Sex* | 75.59 | < 0.001 | 4985.67 | < 0.001 | 0.001 | 0.975 |
| *Type* × *Sex* | 50.99 | < 0.001 | 720.34 | < 0.001 | 27.70 | < 0.001 |

**Figure 1.**
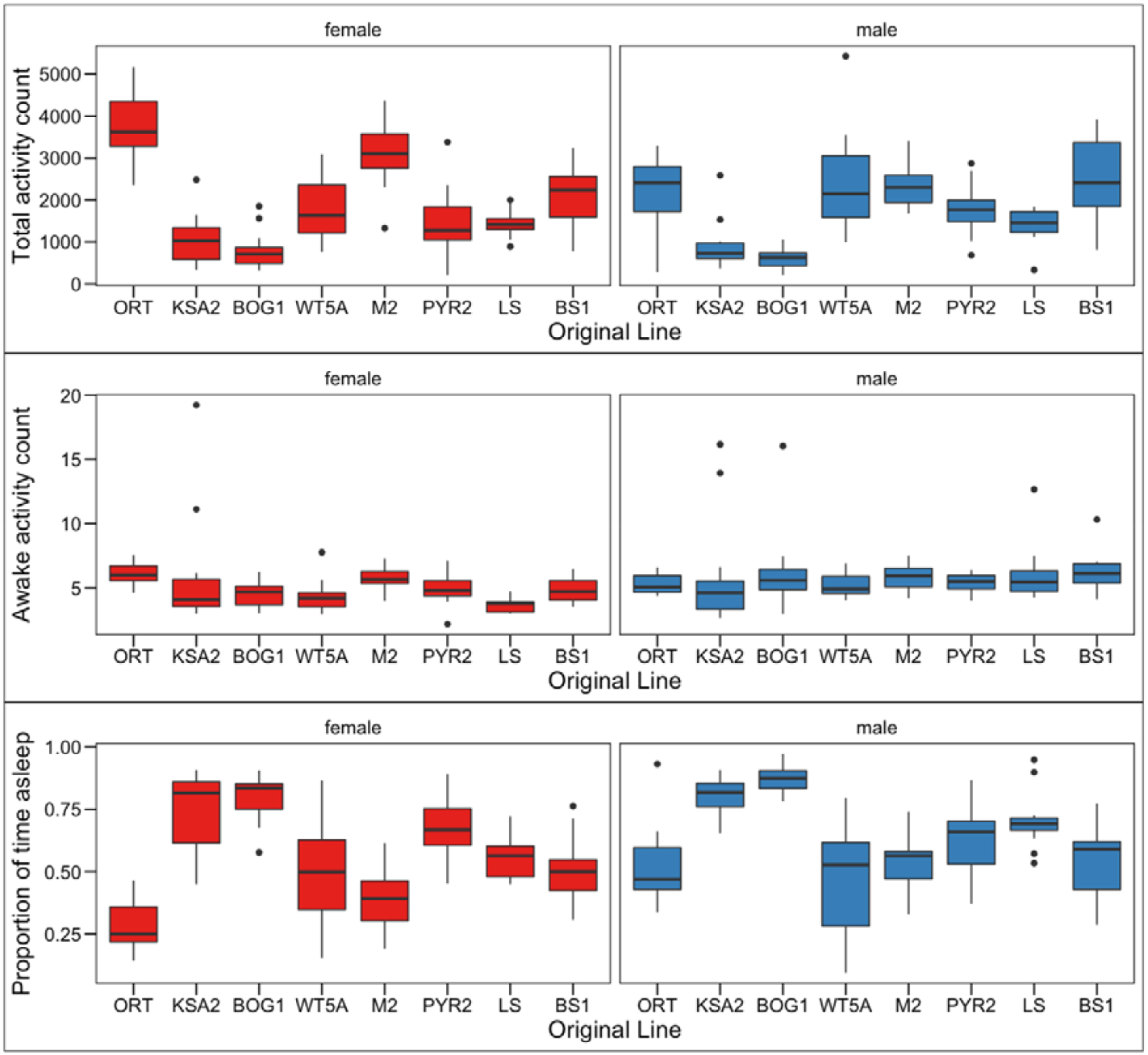
Locomotor activity and sleep in females (red) and males (blue) of the coevolved fly lines. **A)** total number of activity events recorded over three days. **B)** the total number of activity events recorded when the flies were awake. **C)** The proportion of time that flies were determined to be asleep, defined as 5 min of inactivity, See Fig S1 for individual actograms and Table 1 for details of each line. See Table 2 for outputs of statistical models.

### Mitochondrial genome affects activity and sleep patterns of the cybrid lines

By introgressing the eight mtDNA variants onto a common nuclear background (ORT), we were able to evaluate how much of the variance in locomotor activity was affected by the mtDNA and newly created mito-nuclear combinations. In general, comparing total activity counts showed that as observed with co-evolved lines, females of each cybrid line were significantly more active than cybrid males, and the extent of this variation was mtDNA specific (Table 2 ‘Line x Sex’ effect; Figure 2A). Males were found to sleep for a significantly larger proportion of time, although unlike the co-evolved lines, the female cybrids were generally more active when awake when compared to males (Figure 2B-C). Both of these differences were also mediated by mtDNA variation (Table 2 ‘Line x Sex’ effect; Figure 2B-C). Therefore, when the effects of individual mitochondrial genomes are isolated on a common nuclear background females are both more active while awake and also spend less time sleeping than males.

**Figure 2.**
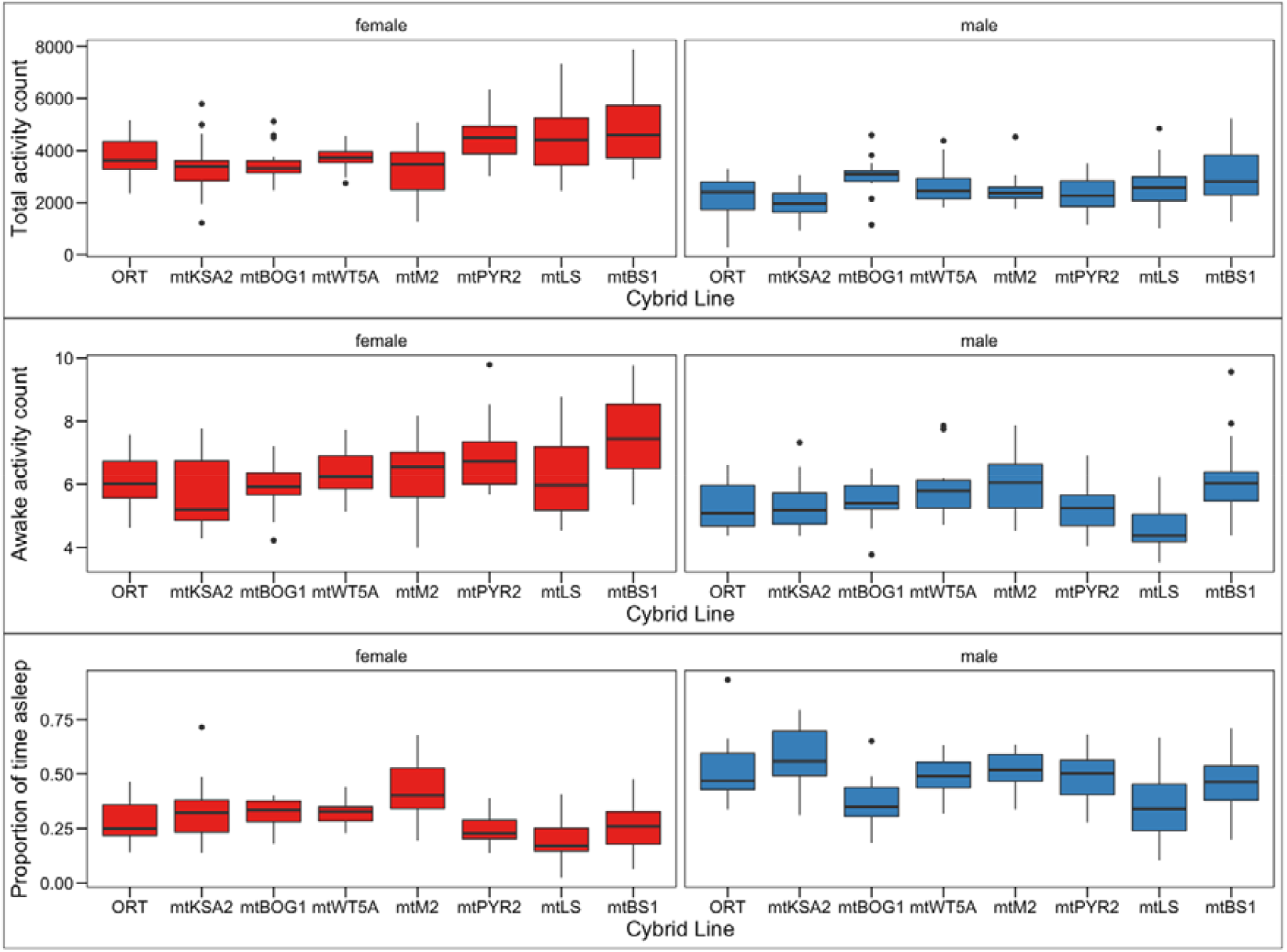
Mitochondrial haplotype effect. Locomotor activity and sleep in cybrid females and males. Each mtDNA variant was introgressed onto the ORT nuclear background (ORT also included here to allow direct comparisons). **A**) total number of activity events recorded over three days. **B**) the total number of activity events recorded when flies were not asleep. **C)** The proportion of time that flies were determined to be asleep, defined as 5 min of inactivity, See Fig S2 for individual actograms and Table 1 for details of each mitochondrial haplotype. See Table 2 for outputs of statistical models.

### Haplogroup specific mtDNA variation can be seen in the activities of cybrid females

Based on the mtDNA coding region variation the eight mtDNA variants form two distinct haplogroups with a set of few common replacement variants present in haplogroup I in OXPHOS complexes I (*ND1;* V190M, *ND2;* I277L, *ND5;* M502I) and V (*ATP6;* S538P and M559V) (Salminen et al. 2017). Haplogroup I contains the haplotypes mtORT, mtKSA2, mtBOG1, mtWT5A and mtM2, as the haplogroup II contains the haplotypes mtPYR2, mtLS and mtBS1 (Table 1). Haplogroup division did not cause clear differences in the sleep-wake activities when the haplogroup specific mtDNA variants were present in their coevolved nuclear backgrounds and the sex differences that were observed earlier with the coevolved lines were decreased (Table 3, Figure 3. upper row). However, when the haplogroup I and II mtDNA variants were placed on a novel common nuclear background in the cybrid lines, we observed that haplogroup II females we more active than haplogroup I females (Table 3, Figure 3, lower row A). Haplogroup II females were also more active when awake and slept for a smaller proportion of time than haplogroup I females (Table 3, Figure 3, lower row B and C).

**Table 3.**
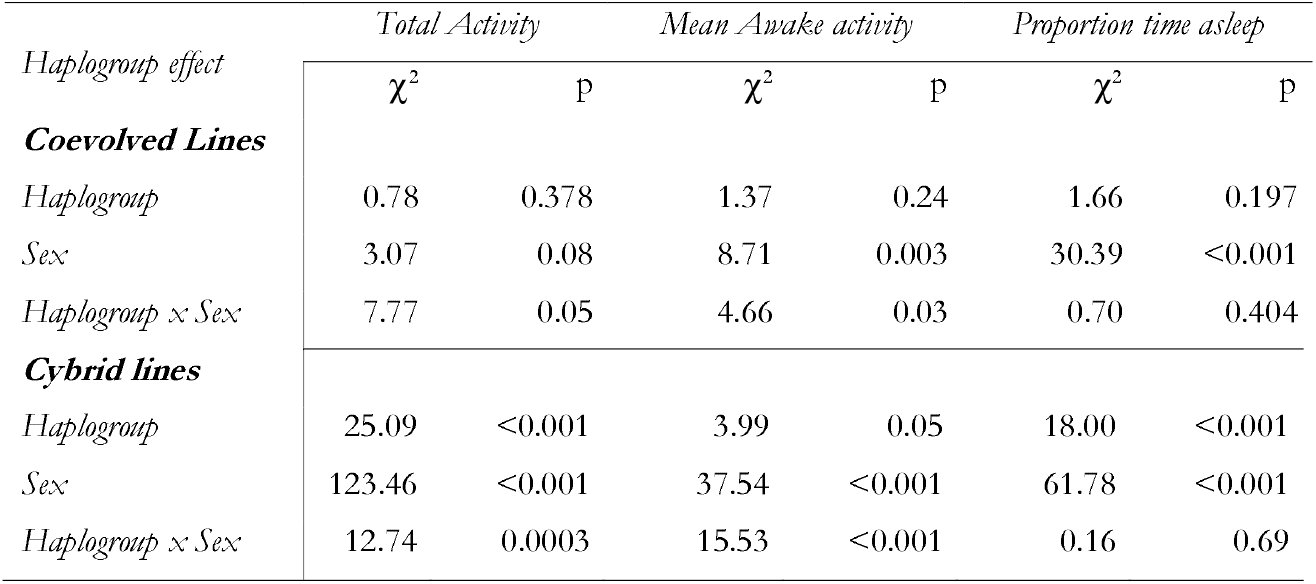
Summaries of test statistics for fixed effects in testing the effect of haplogroup and sex on activity and sleep in coevolved and cybrid lines. See Figure 3 and methods for model details.

**Figure 3.**
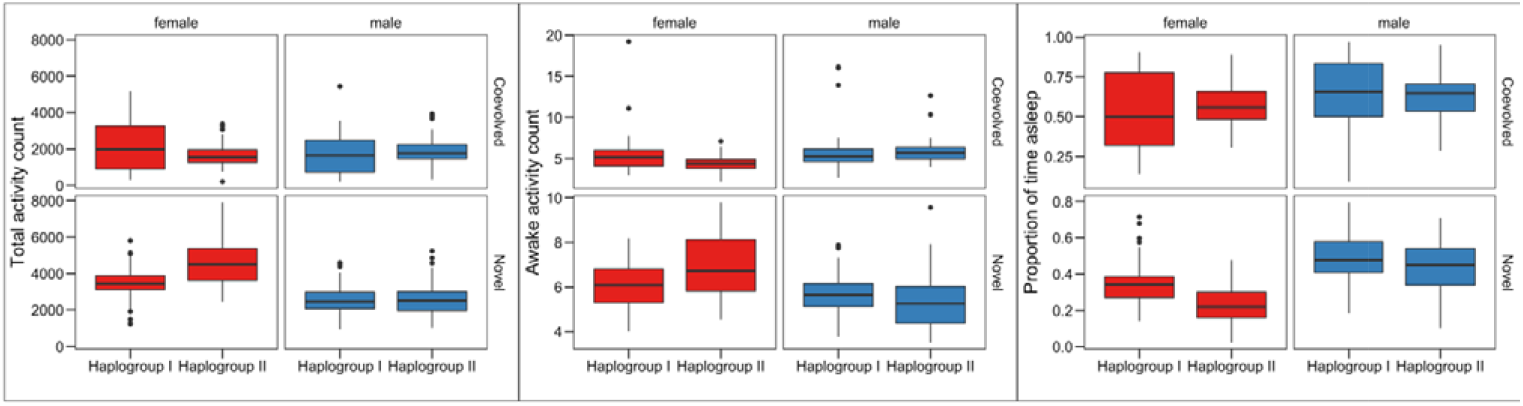
Haplogroup effect. Locomotor activities and sleep in coevolved and cybrid females and males based on the mtDNA haplogroup division. A) Total activity counts over three days. B) Activity of the flies when awake. C) The proportion of time the flies spent sleeping.

### Breaking up co-evolved mito-nuclear combinations affects activity and sleep patterns

As naturally occurring mito-nuclear genome combinations have co-adapted locally over time, we expected that disruption of these combinations could result in maladaptive effects. We also expected that these effects might be more severe, or more variable in males, as postulated under the mother’s curse hypothesis (65-68). Both male and female cybrid flies harbouring novel mito-nuclear combinations were significantly more active than co-evolved flies, although the extent of this variation differed between sexes and was more prominent in females (Table 2 ‘Line x Sex’ effect; Figure 4A). This shift is partially due to the Oregon RT nuclear background, as shown it Figure 1A ORT females had the highest total activity counts when compared to other co-evolved lines. Part of the increase in total activity in cybrids was driven by increase in the awake activity level, though this was the case only in females (Table 2 ‘Line x Sex’ effect; Figure 4B). However, the largest driver of the increased total activity in both male and female cybrid flies is that these spent a significantly lower proportion of their time asleep compared to the co-evolved lines, the extent of this difference was slightly larger in females (Table 2 ‘Line x Sex’ effect; Figure 3C) and again, this is largely due to ORT nuclear background normalizing the variation of the cybrid lines when compared the co-evolved lines (Fig. 1C, ORT females spent the least time asleep).

**Figure 4.**
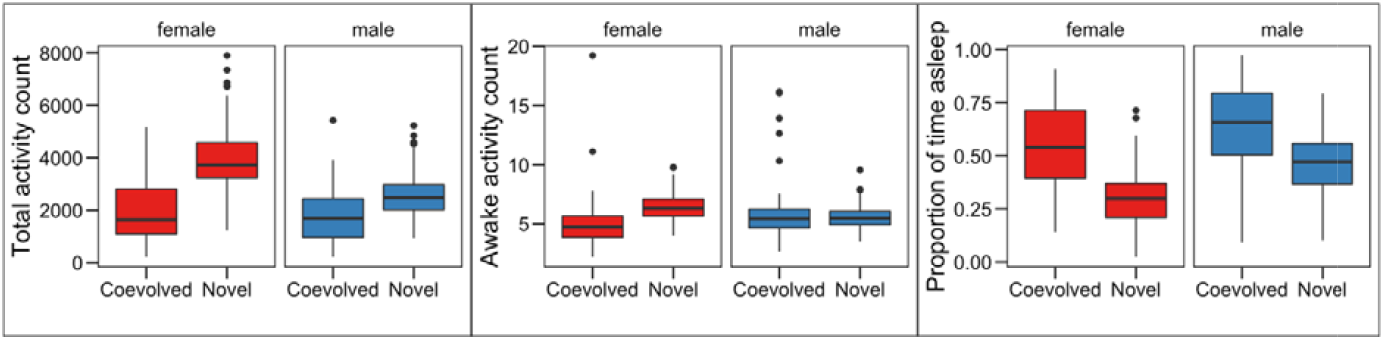
A comparison of Coevolved and Novel Mito-nuclear associations. A) total number of activity events recorded over 3 days. B) the total number of activity events recorded when flies were not asleep. C) The proportion of time that flies were determined to be asleep, defined as 5 min of inactivity, See Table 2 for outputs of statistical models.

### Cybrid activity levels are positively correlated with mtDNA copy number

We utilized an earlier published mtDNA copy number data collected from the same co-evolved and cybrid lines as studied in this manuscript (37). Salminen *et al*. (2017) measured mtDNA copy number separately from females and males from various timepoints after eclosion. For our purposes we selected 3-days post eclosion mtDNA copy number data as this was the closest match with the age of most the flies used in our activity measurements. Our aim was to test the correlation between mtDNA copy number and activity levels of the co-evolved and cybrid females and males. When mtDNA copy number was correlated with the total activity counts of the cybrid lines, we saw a significant positive correlation in cybrid males and similar but non-significant trend in cybrid females, indicating that with increasing mtDNA copy number levels the total activity of the flies also increased (Fig. 5A, right panel). We also saw a weak positive trend between awake activity levels and mtDNA copy number (Fig. 5B, right panel) and a weak negative trend between mtDNA copy number and sleep (Fig. 5C). In co-evolved lines we did not detect evidence of a significant correlation between mtDNA copy number and activity or sleep (Figure 5A-C, left panel).

**Figure 5.**
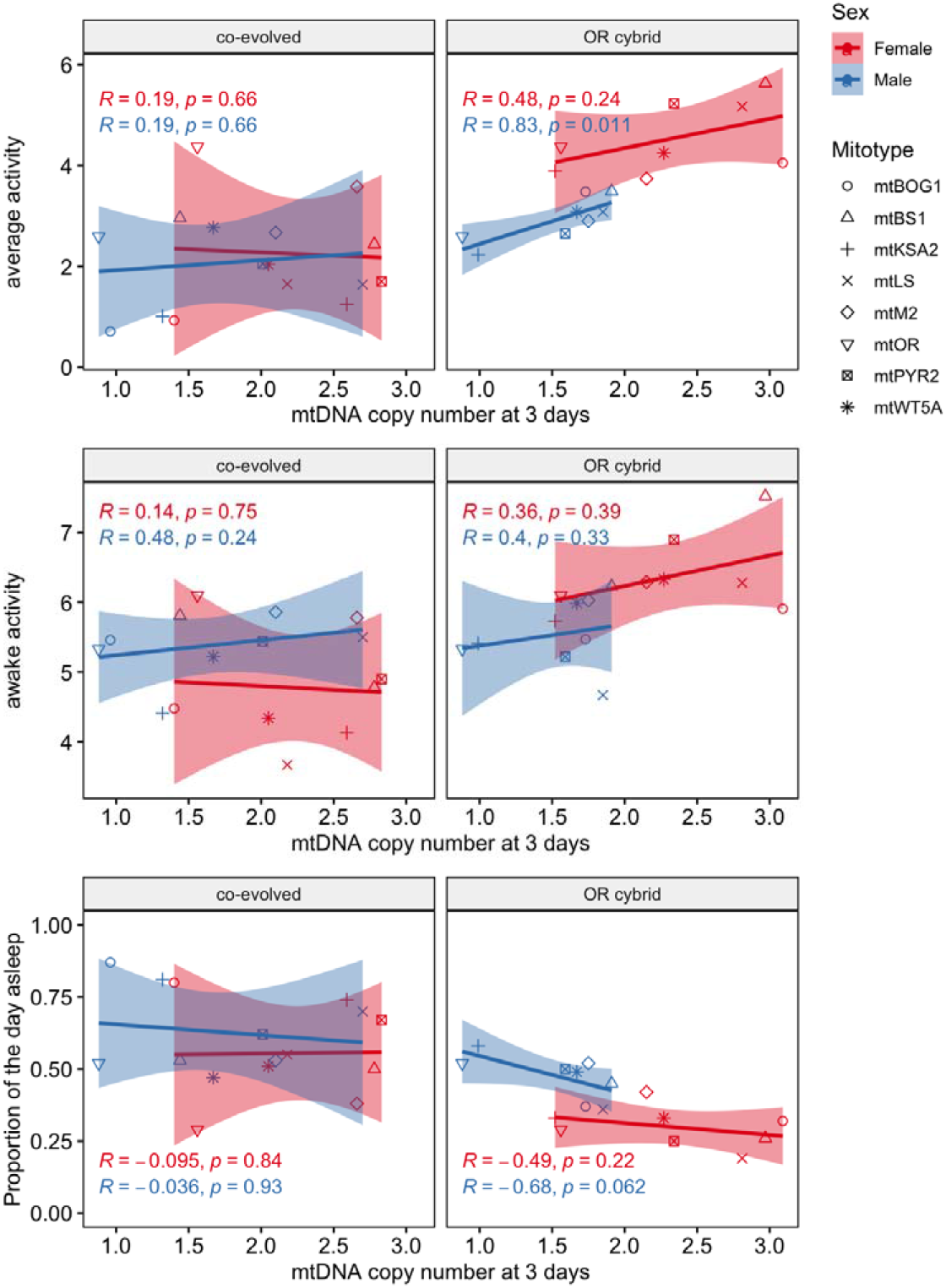
mtDNA copy number correlations between measured traits. Spearman’s rank correlations between average mtDNA copy number and averages of a) total number of activity events, b) total number of activity events when the flies were awake and c) proportion of time the flies were determined to be asleep measured from co-evolved (left panel) and cybrid (right panel) females and males.

## Discussion

Mitochondrial dysfunction commonly manifests in phenotypes related to energy-demanding tissues, such as muscles, and movement disorders are a major feature of mitochondrial diseases (53). *Drosophila* can be employed as a model for as movement disorders, and its locomotive behaviours include larval crawling, adult walking, climbing and flight. These behaviours can be used to evaluate muscle behaviour as a proxy of e.g. mitochondrial performance. Furthermore, periods when the are immobile for longer than five-minutes, are typically considered sleeping (39). Sleep deprivation has been shown to be connected to mitochondrial bioenergetics and to cause mitochondrial dysfunction in *Drosophila* (54). However, investigation of sleep as a consequence of mitochondrial dysfunction appears to be an understudied aspect of mitochondrial disease (55). It is also unclear how retrograde signalling from mitochondria to nucleus affects sleep-wake cycles, and especially the role of naturally occurring mtDNA variation in these traits.

Studying the effect of mtDNA variation is challenging as it is necessary to disentangle the effects of the mitochondrial genome from the effect of the nuclear genome. This is feasible with the *Drosophila* cybrid model and here we have focused on studying the effect of mtDNA variation and mito-nuclear interactions on locomotion activity and sleep. We addressed three questions by separately quantifying the contributions of the mitochondrial and nuclear genomes to activity and sleep phenotypes. First, we asked how variation in both the nDNA and mtDNA affects sleep and activity. Second, by isolating mtDNA variants on a common nuclear background we investigated how variation in mtDNA affects sleep and activity. Finally, we assessed how breaking up co-evolved mito-nuclear genetic interactions affected the sleep and activity.

Locomotor activity and the proportion of time spent asleep were first measured from eight wild type Drosophila strains with co-evolved mito-nuclear combination adapted to their local environment. These strains showed variation in their sleep-wake patterns and in general the females were found to be more active overall; males exhibited a higher waking activity but slept for a larger proportion of time. We next aimed to test if mtDNA contributed to this variation between lines. The eight studied mtDNA genomes can be subdivided into two haplogroups based on their genetic variation (37). These haplogroups differed in the amino acid replacements present in OXPHOS complex I genes *ND1, ND2* and *ND5*, as well as two variants from complex V gene *ATP6* (37). Based on these common replacement variants, haplogroup II contained strains of European origin, whereas haplogroup I contained strains from different continents (37). Interestingly, haplogroup II females were shown to have higher total activity levels and spent less time in sleeping when compared to haplogroup I females. The same was not observed with males. It is unclear how these patterns might have arisen in nature, however further experimentation is required to examine of these patterns are a result of local adaptation or just due to drift.

As the eight cybrids also possess mitochondrial haplotype-specific mutations, we examined the effect of these to locomotion and sleep, when introgressed into common nuclear background in the cybrid lines. In general, we were able to see differences in the amounts of activity and sleep, in both females and males, brought upon by mtDNA variation. Cybrid females were more active than males, sleep less and have higher waking activity. It is difficult to say if specific mtDNA mutations are causing the seen variation, as most of these mtDNA genomes contain more than one source of variation, i.e replacement variants in the protein coding genes, synonymous SNPs, indels in the tRNA and rRNA genes and also length variation in the non-coding A+T region (37). Mitovariant mtM2, the only variant that is originally from Australia, contains the most unique replacement variants when compared to the other mitovariants (Table 1). mtM2 was also one of the lowest activity strains, and females especially spent more time sleeping when compared to the other cybrids. This might be due to altered interactions between mitochondrial and nuclear gene products (56). M2 flies which have the original nuclear background actually appear to exhibit relatively high activity and low amount of sleep. Disruption of naturally occurring mito-nuclear combinations can result in interruption of precise interactions, leading to decreased mito-nuclear cooperation and a reduction in fitness (36).

mtKSA2 is the only African variant, and it possesses two unique amino acid replacement variants in OXPHOS cIII (*CYTB*) and cIV (*COIII*) when compared to other cybrids (37). mtKSA2 cybrids males exhibited the highest proportion of time spent sleeping. Also, the overall activity counts were among the lowest in mtKSA2 females and males, and the mtKSA2 cybrids females also had the lowest awake activity counts. The mtDNA variation studied here is maternally inherited and its effects can be multisystemic, affecting each tissue in Drosophila. However, there are cases where several sporadic missense and nonsense *CYTB* mutations in muscles have been shown to cause complex III deficiency and exercise intolerance in humans (57). *CYTB* mutation in mtKSA2 may partially explain the lower activity rates when compared to other strains.

mtKSA2 is also associated with low mtDNA copy number in ORT nuclear background when compared to other cybrid lines (37). Since mtDNA copy number has been shown to be associated with other measures of fitness in Drosophila including fertility and longevity (58) and with development time and weight (37) of the same co-evolved and cybrid lines as studied here, we hypothesised that it might also be associated with variation in activity levels. mtDNA copy number is sexually dimorphic in Drosophila, with females of most strains tending to exhibit a higher copy number than males (37, 58), and is also affected by the age of the flies, sex specifically (37). Sleep and activity are also shown to be sexually dimorphic traits in Drosophila (39, 59). In our study we saw that in both co-evolved and cybrid strains the females were more active overall and slept less, as shown also with previous *Drosophila* sleep and activity research (60). These factors motivated us to analyse whether the earlier measured mtDNA copy number levels of the co-evolved and cybrid females and males correlate with the traits measured in this manuscript. We expected that higher activity and lower sleep exhibited by female flies may be at least partly due to their higher mtDNA copy number. This was the case with the cybrid lines, but was not seen with the co-evolved lines. The strongest positive correlation was found in cybrid males between mtDNA copy number and mean activity levels. This data shows that in controlled nuclear background the mtDNA effect on both mtDNA copy number and activity levels have a stronger penetrance compared to co-evolved lines, where nuclear variation might mask the effect of mtDNA variation on copy number and activity. In addition, the low mtDNA copy number exhibited by nOR mtKSA2 among the cybrids is likely responsible for the strain’s reduced activity in some degree.

Comparison of the co-evolved mito-nuclear strains with the novel mito-nuclear strains allowed investigation into the effects of mito-nuclear epistatic interactions and the effects of breaking up possible co-evolved mito-nuclear gene complexes on organismal fitness. Previous studies have demonstrated that disrupting these interactions can lead to decreased OXPHOS function (37, 61). Specifically, cytochrome c oxidase activity has been shown to be reduced when an mtDNA haplotype is combined with a more distant nuclear background rather than its original nuclear background in copepod *Tigriopus californicus* (62). Mito-nuclear incompatibilities can lead to reduction in fitness (63-65), and maintaining normal levels of activity is beneficial to fitness to allow, for example foraging and mating to take place (59). Short sleep duration in Drosophila has previously been shown to be associated with poor memory (66), and reduced longevity (67, 68). Therefore, it is likely that the altered sleep and activity patterns exhibited by cybrid strains are likely to equate to a reduction in fitness, supporting the theory of mito-nuclear co-adaptation.

Natural selection is blind to deleterious mitochondrial phenotypes which manifest in males as mtDNA is maternally inherited (69-71). This can indicate that variation which persists in mtDNA is more likely to cause greater phenotypic divergence within males and more likely to cause deleterious effects in males than in females (69, 71). Although studies exist which provide evidence in support of this theory, known as the mother’s curse (33, 72), based on our results, it does not appear that the effect on sleep or activity in males is more prominent than in females when breaking up the co-evolved combinations, and does not provide evidence in support of the mother’s curse.

Drosophila sleep and activity research will enable further understanding of the causes of abnormal sleep and activity patterns in humans and the role of mitochondrial variation in these traits. Further studies of mito-nuclear interactions and mtDNA variation are essential in recognizing and eventually preventing mito-nuclear mismatches that might occur during mitochondrial replacement theory.

## Acknowledgements

We thank Angela Reid and Lucinda Rowe and the Ashworth Media team for fly medium preparation. Lucy Anderson was supported by the Genetics Honours Program, School of Biological Sciences, University of Edinburgh. This work was partially supported by a Chancellor’s Fellowship (University of Edinburgh), a Branco Weiss Fellowship (Society in Science – Zürich, https://brancoweissfellowship.org/) and a Leverhulme Trust Research Grant (RPG-2018-369), all awarded to Pedro Vale.

## Supplementary Figures and tables

Fig S1 – Actograms for original lines with coevolved nuclear and mitochondrial genomes

Fig S2 – Actograms for cybird lines with novel nuclear and mitochondrial combinations

**Fig S1.**
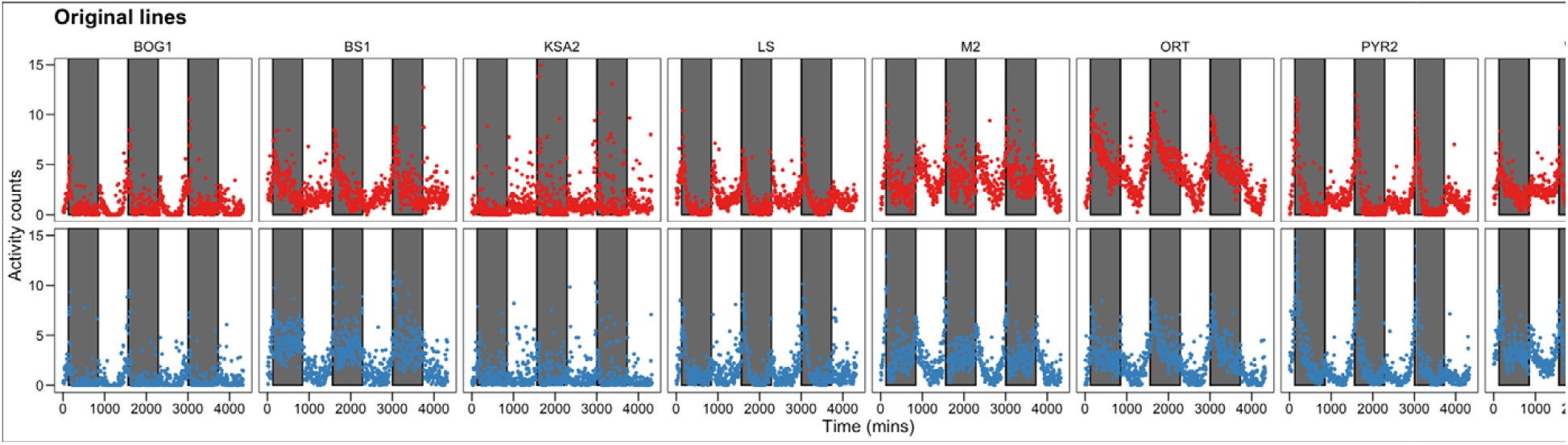
Drosophila Activity Monitor (DAM) Activity counts (unique breaks of the infra-red beam) measured for each original fly line over a period of three days. Females are shown in the top panel in red; males in the bottom panel in blue. Within each plot, the white vertical bars indicate periods of light, while the dark vertical bars indicate periods of dark when lights are turned off within the incubator. Details about each line can be found in Table 1. Summary statistics for the activity patters can be found in Figure 1.

**Fig S2.**
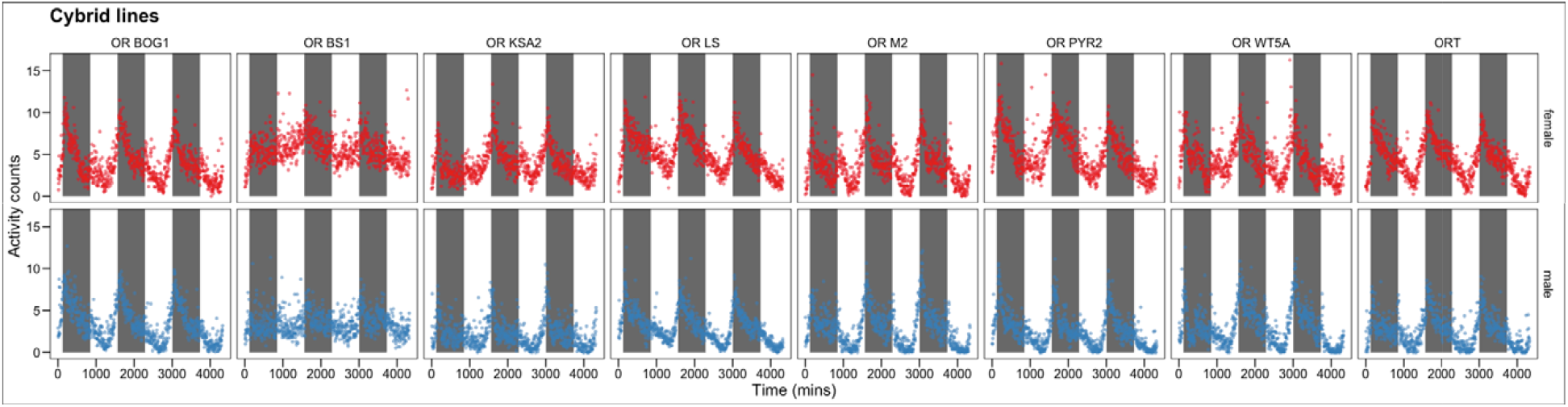
Drosophila Activity Monitor (DAM) Activity counts over a period of three days (unique breaks of the infra-red beam) measured for each cyrid fly line, where each mitochondrial haplotype has been introgressed onto the OR nuclear background. Females are shown in the top panel in red; males in the bottom panel in blue. Within each plot, the white vertical bars indicate periods of light, while the dark vertical bars indicate periods of dark when lights are turned off within the incubator. Details about each line can be found in Table 1. Summary statistics for the activity patters can be found in Figure 2.

## Notes

https://doi.org/10.5281/zenodo.5573904

### Competing Interest Statement

The authors have declared no competing interest.

